# Ubiquitin transfer by a RING E3 ligase occurs from a closed E2~Ub conformation

**DOI:** 10.1101/848788

**Authors:** Emma Branigan, J. Carlos Penedo, Ronald T. Hay

## Abstract

Ubiquitination is a eukaryotic post-translational modification that modulates a host of cellular processes^1^. Modification is mediated by an E1 activating enzyme (E1), an E2 conjugating enzyme (E2) and an E3 ligase (E3). The E1 catalyses formation of a highly reactive thioester linked conjugate between ubiquitin and E2 (E2~Ub)^2^. The largest class of ubiquitin E3 ligases, which is represented by RING E3s, bind both substrate and E2~Ub and facilitate transfer of ubiquitin from the E2 to substrate. Based on extensive structural analysis^3–5^ it has been proposed that RING E3s prime the E2~Ub conjugate for catalysis by locking it into a “closed” conformation where ubiquitin is folded back onto the E2 exposing the restrained thioester bond to attack by a substrate nucleophile. However the proposal that the RING dependent closed conformation of E2~Ub represents the active form that mediates ubiquitin transfer is a model that has yet to be experimentally tested. Here we use single molecule Förster Resonance Energy Transfer (smFRET) to test this hypothesis and demonstrate that ubiquitin is transferred from the closed conformation during an E3 catalysed reaction. Using Ubc13 as an E2, we designed a FRET labelled E2~Ub conjugate, which distinguishes between closed and alternative conformations. Firstly, we defined the high FRET state as the closed conformation using a stable isopeptide linked E2~Ub conjugate, while the low FRET state represents more open conformations. Secondly, we developed a real-time smFRET assay to monitor RING E3 catalysed ubiquitination with a thioester linked E2~Ub conjugate and determined the catalytically active conformation. Our results demonstrate that the reaction proceeds from the high FRET or closed conformation and confirm the hypothesis that the closed conformation is the active form of the conjugate. These findings are not only relevant to RING E3 catalysed ubiquitination but are also broadly applicable to E3 mediated ligation of other ubiquitin-like proteins (Ubls) to substrates.

## Results and Discussion

Ubiquitin in an E2~Ub conjugate can exist in multiple conformations including a “closed” form in which ubiquitin is folded back onto the E2. E3 ligases for ubiquitin and Ubl proteins appear to trap the E2~Ub/Ubl conjugate in this conformation^3–11^ (Fig. 1a). We have previously crystallized Ubc13~Ub in complex with the RNF4 RING domain dimer without the Ubc13 binding partner, UEV, in the same closed conformation^7^ (Fig. 1b) even although RNF4 fails to transfer ubiquitin from the Ubc13~Ub conjugate in the absence of UEV (Fig. 1c). Here we found that RNF4 has a higher binding affinity for the Ubc13~Ub conjugate when in complex with UEV (Fig. 1d) suggesting both UEV and RNF4 are required to stabilize the active conformation of the conjugate. We therefore decided to use smFRET to interrogate the structural dynamics of the Ubc13~Ub conjugate in response to RNF4 and UEV binding and RING E3 catalysed ubiquitin transfer.

**Figure 1.**
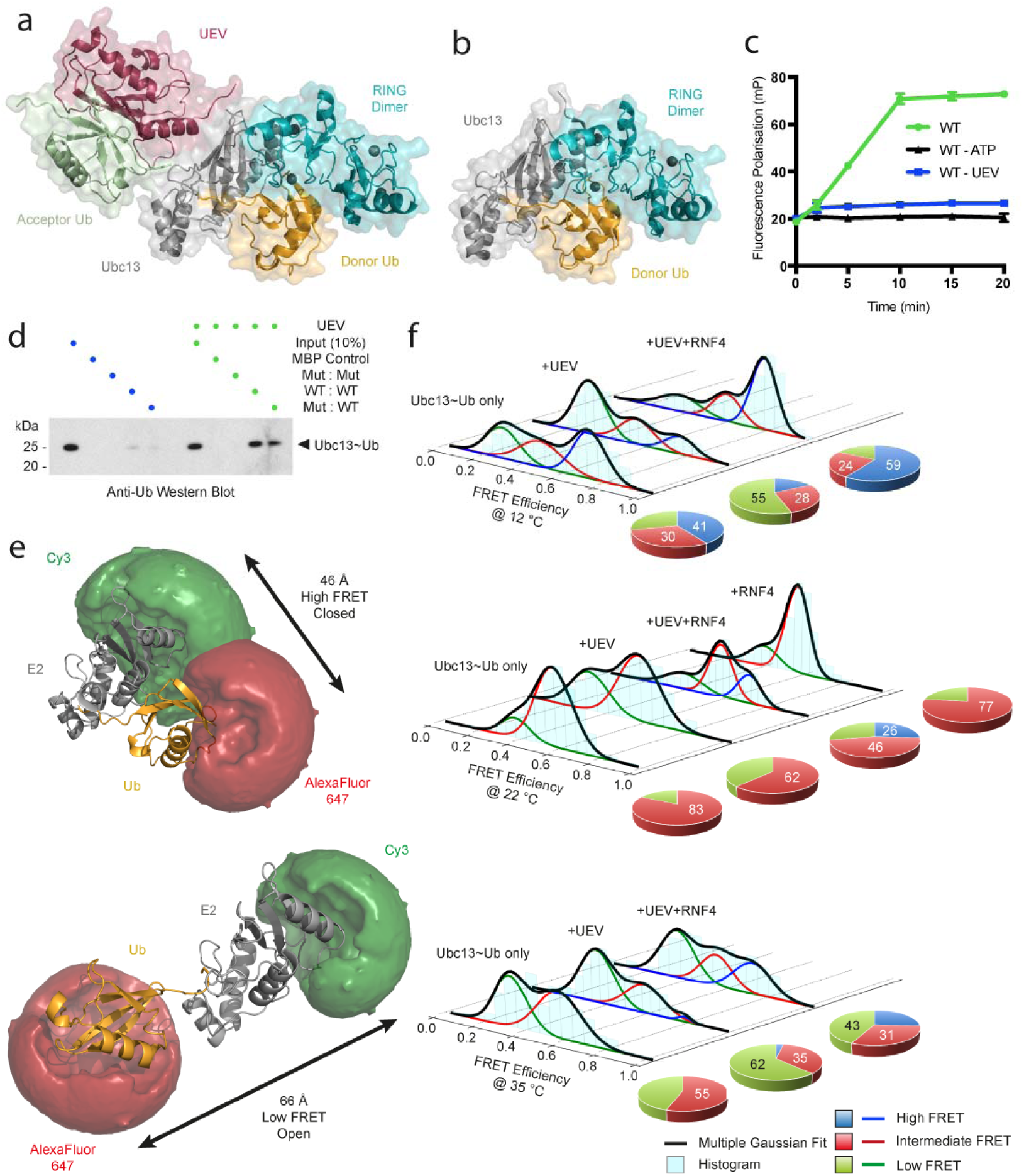
UEV and the RNF4 RING domain dimer are required to stabilize the closed conformation of the E2~Ub conjugate. **a**, Structural model of the RNF4 RING domain dimer in complex with the Ubc13~Ub conjugate, UEV and acceptor ubiquitin (PDB 5AIT). **b**, Crystal structure of the RNF4 RING domain dimer in complex with the Ubc13~Ub conjugate alone (PDB 5AIU). For **a** and **b**, binding of only one Ubc13~Ub conjugate is displayed for clarity. **c**, Fluorescence polarization ubiquitination assay showing the effect of UEV on K63-linked polyubiquitin chain formation, represented as mean ± s.d. of triplicate measurements. Error bars are omitted when the error is smaller than the data point. **d**, RNF4 RING domain dimer pull-down experiment showing the effect of UEV on Ubc13~Ub conjugate binding, analysed by Western blot. WT denotes a wild-type RING domain, while Mut denotes a RING domain containing E2~Ub binding site mutations (M140A and R181A). **e**, Modelled accessible volume of FRET dyes and average distance measurement for E2~Ub conjugate closed and open conformations. **f**, smFRET histograms showing the isopeptide linked E2~Ub conjugate conformation alone and in complex with UEV and the RNF4 RING domain dimer at three different temperatures. The E2~Ub conjugate conformation in the presence of the RNF4 RING domain dimer only is shown at 22 °C. Pie charts to the right of each histogram show the percentage contribution of each FRET state to the overall population.

Single molecule fluorescence approaches have been applied to the ubiquitin system^12–14^; but not to follow RING catalysed transfer of ubiquitin from E2 to substrate. We thus designed a Ubc13~Ub conjugate labelled with the Cy3B-AlexaFluor 647 FRET pair (R_0_ = 60 Å) where proximity of the FRET labels in the closed conformation yields a high FRET efficiency, while the large distance between the labels in the more open conformation results in a low FRET efficiency (Fig. 1e, Extended Data Fig. 1a, b). We used a stable isopeptide linked Ubc13~Ub conjugate^3,7^ to first determine the FRET state of the conjugate in solution and the effect that UEV and the RNF4 RING domain dimer has on its conformation. The resulting conjugate bound with similar efficiency to the RNF4 RING domain dimer as the untagged and unlabelled conjugate (Fig. 1d, Extended Data Fig. 1c).

The Ubc13~Ub conjugate alone has access to a wide range of FRET states at low temperature (12 °C) and exhibits a slight preference (41%) for a high FRET state with a FRET efficiency value (E_FRET_) of 0.71, indicating that the closed conformation is favoured (Fig. 1f, Extended Data Fig. 2a). At higher temperatures (22 and 35 °C), an intermediate FRET state (E_FRET_~0.57) and a low FRET state (E_FRET_~0.36) are favoured and no high FRET state was observed (Fig. 1f, Extended Data Fig. 2b, c, Extended Data Fig. 3a). In the presence of UEV at all temperatures, the low FRET population is stabilized while the intermediate and high FRET states are destabilized, indicating a more open conformation of the conjugate in complex with UEV (Fig. 1f, Extended Data Fig. 2a-c). The addition of the RNF4 RING domain dimer to the Ubc13~Ub conjugate in complex with UEV at all temperatures, captures the high FRET state, consistent with stabilization of a closed or active conformation of the Ubc13~Ub conjugate (Fig. 1f, Extended Data Fig. 2a-c, Extended Data Fig. 3b). In the absence of UEV, the RNF4 RING domain dimer does not stabilize the high FRET state (Fig. 1f, Extended Data Fig. 2b), showing that a preassembled UEV/Ubc13~Ub complex is required for the RING dimer to capture the closed conformation. Analysis of mutations that disrupt UEV-E2 and RNF4-E2~Ub interactions confirmed that changes in FRET were due to specific binding of UEV and RNF4 RING domain dimer to the Ubc13~Ub conjugate (Extended Data Fig. 4a-c). Increased temperature is anticipated to destabilize the high FRET state or closed conformation due to breakage of the hydrogen-bonding network in the active site groove that is required for adopting the closed conformation and drives specificity for catalysis in the active site^3^. As a result the C-terminal tail of ubiquitin will sit outside the active site groove similar to a previous NMR model^15^ leading to a sub-optimal conformation corresponding to the intermediate FRET state (Figure 1f). However, it is likely that the hydrophobic interaction between ubiquitin and Ubc13 in the E2~Ub conjugate, is still engaged in the intermediate FRET state.

To test this directly and confirm that the high FRET state represents the closed conformation we used the I44A mutation in ubiquitin (Fig. 2a) that is inactive in ubiquitination in vitro (Fig. 2b) and disrupts the closed conformation by blocking the interaction between the hydrophobic patch of ubiquitin and E2. This mutation stabilizes the low FRET state indicating a more open conformation and the conjugate cannot access the high FRET state or closed conformation, even in the presence of UEV and the RNF4 RING domain dimer (Fig. 2c, Extended Data Fig. 5a). An L106A mutation in Ubc13 that is part of the same hydrophobic interface has a similar, but less pronounced, effect on the conformation of the Ubc13~Ub conjugate (Fig. 2 a, b, d, Extended Data Fig. 5b). In addition to destabilizing the high FRET state, these mutations also destabilize the intermediate FRET state, indicating that the hydrophobic interaction maintains both the intermediate and high FRET states, and when disrupted a more open conformation or low FRET state is favoured. An S32A mutation in UEV^16^ that perturbs binding of the acceptor ubiquitin (Fig. 2a), was defective in RNF4 dependent substrate ubiquitination (Fig 2b), but retained the ability to activate the Ubc13~Ub conjugate in an RNF4 dependent lysine discharge assay^7^. To determine if the S32A mutation in UEV could co-operate with RNF4 to restrain the Ubc13~Ub conjugate in the closed conformation we carried out smFRET analysis of the wild type and S32A versions of UEV. This revealed that the conformation of the Ubc13~Ub conjugates were indistinguishable and indicated that stabilization of the closed conformation was independent of acceptor ubiquitin binding to UEV (Fig. 2e). These data also indicate that none of the FRET states observed are a consequence of ubiquitin binding to UEV.

**Figure 2.**
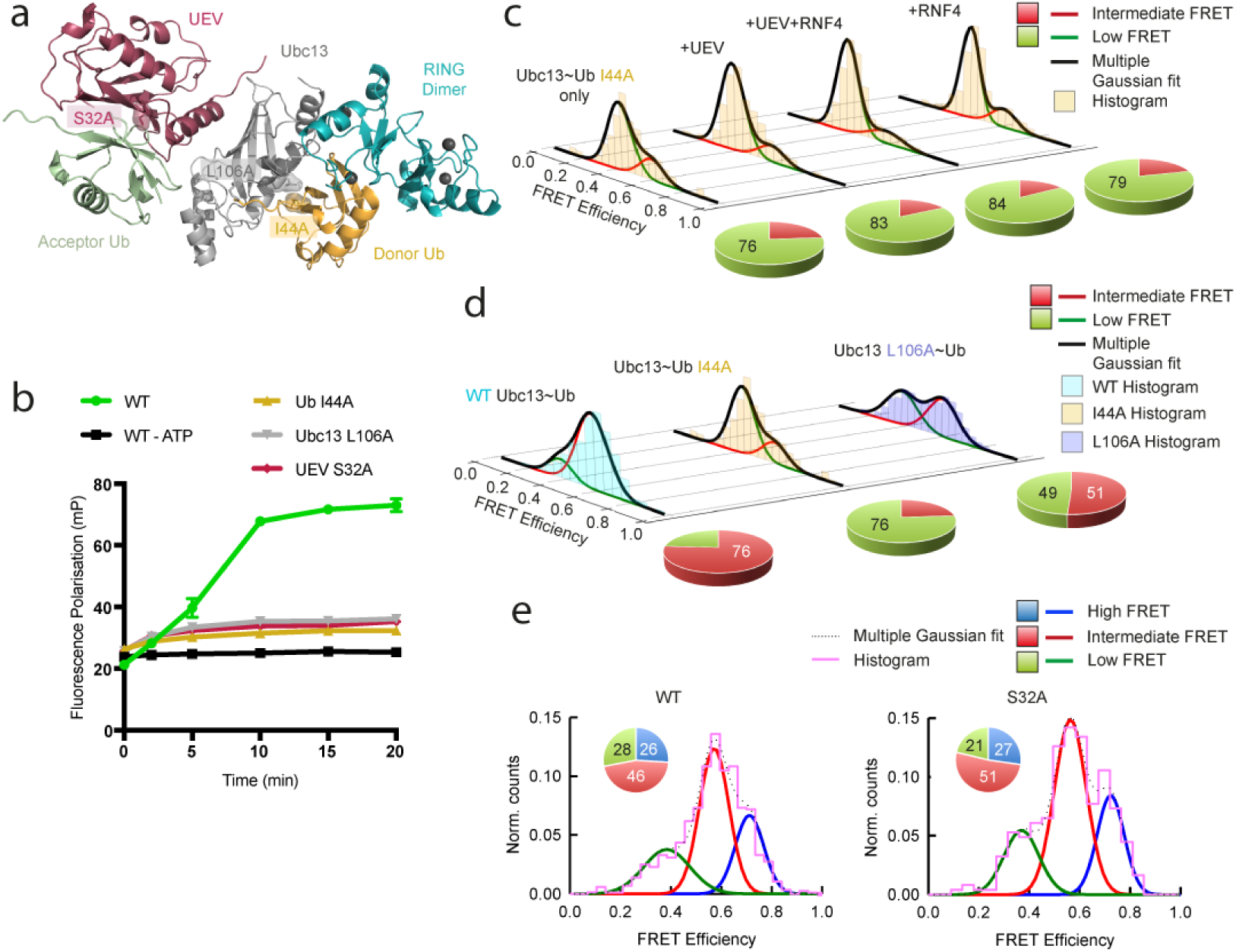
Effects of complex mutagenesis on the conformational state of the E2~Ub conjugate. **a**, Locations of mutations made within the complex (PDB accession number 5AIT). **b**, Fluorescence polarization ubiquitination assay showing the effect of mutations on K63-linked polyubiquitin chain formation, represented as mean ± s.d. of triplicate measurements. Error bars are omitted when the error is smaller than the data point. **c**, smFRET histograms showing the E2~Ub conjugate conformation containing Ub I44A alone and in complex with UEV and the RNF4 RING domain dimer. **d**, smFRET histograms showing the conformation of the WT E2~Ub conjugate and the E2~Ub conjugate containing either Ub I44A or Ubc13 L106A. For **c** and **d**, pie charts to the right of each histogram highlight the percentage contribution of each FRET state to the overall population. **e**, Normalized smFRET histograms showing conformation of the E2~Ub conjugate in complex with the RNF4 RING domain dimer and either WT UEV (left panel) or UEV S32A (right panel). The histogram is outlined in magenta. Pie chart insets show the percentage contribution of each FRET state to the overall population.

Using the stable isopeptide linked Ubc13~Ub conjugate allowed us to identify the high FRET state as the closed conformation and indicated that both UEV and RNF4 were required to restrain the Ubc13~Ub conjugate in this conformation. However, to directly test the hypothesis that the closed conformation is the active form of the Ubc13~Ub conjugate, requires a method that reports in real-time the FRET state(s) from which ubiquitin transfer to substrate occurs. This necessitated attaching the same FRET dyes at identical positions used for the isopeptide linked conjugate to an unstable thioester linked conjugate (Fig. 3a, Extended Data Fig. 6a, b). The labelled conjugate modified the substrate in a biochemical assay with similar efficiency to the WT conjugate (Extended Data Fig. 7). In the smFRET experiment we immobilized the unstable thioester linked Ubc13~Ub conjugate onto the surface for imaging and injected the reaction components for ubiquitination including UEV, RNF4 and Ub~4xSUMO-2 substrate. We monitored, in real-time, transfer of Cy3B-labelled ubiquitin from the E2~Ub conjugate onto substrate, which diffuses into solution and results in simultaneous loss of both donor and acceptor signals, and therefore, the FRET signal (Fig. 3a). This is evident from the decrease in number of FRET pairs before and after the reaction has occurred (Fig. 3b) and by following the cumulative intensity of fluorescent spots over time with and without RNF4 catalysed ubiquitination (Fig. 3c). Single molecule trajectories demonstrate stable Cy3B and AlexaFluor 647 intensities lasting hundreds of seconds following a UEV only injection versus the rapid simultaneous loss of all fluorescent signals following the ubiquitination reaction (Fig. 3d). Combined analysis of single molecule traces shows a significant portion of E2~Ub conjugates react upon injection of reaction components (~60%), compared to injection of UEV only (~14%) (Fig. 3e). A small portion of the sample contains only Cy3B due to under-labelling with AlexaFluor 647 or prior photobleaching of AlexaFuor 647, however, a reaction was also observed in these conjugates (Fig. 3e).

**Figure 3.**
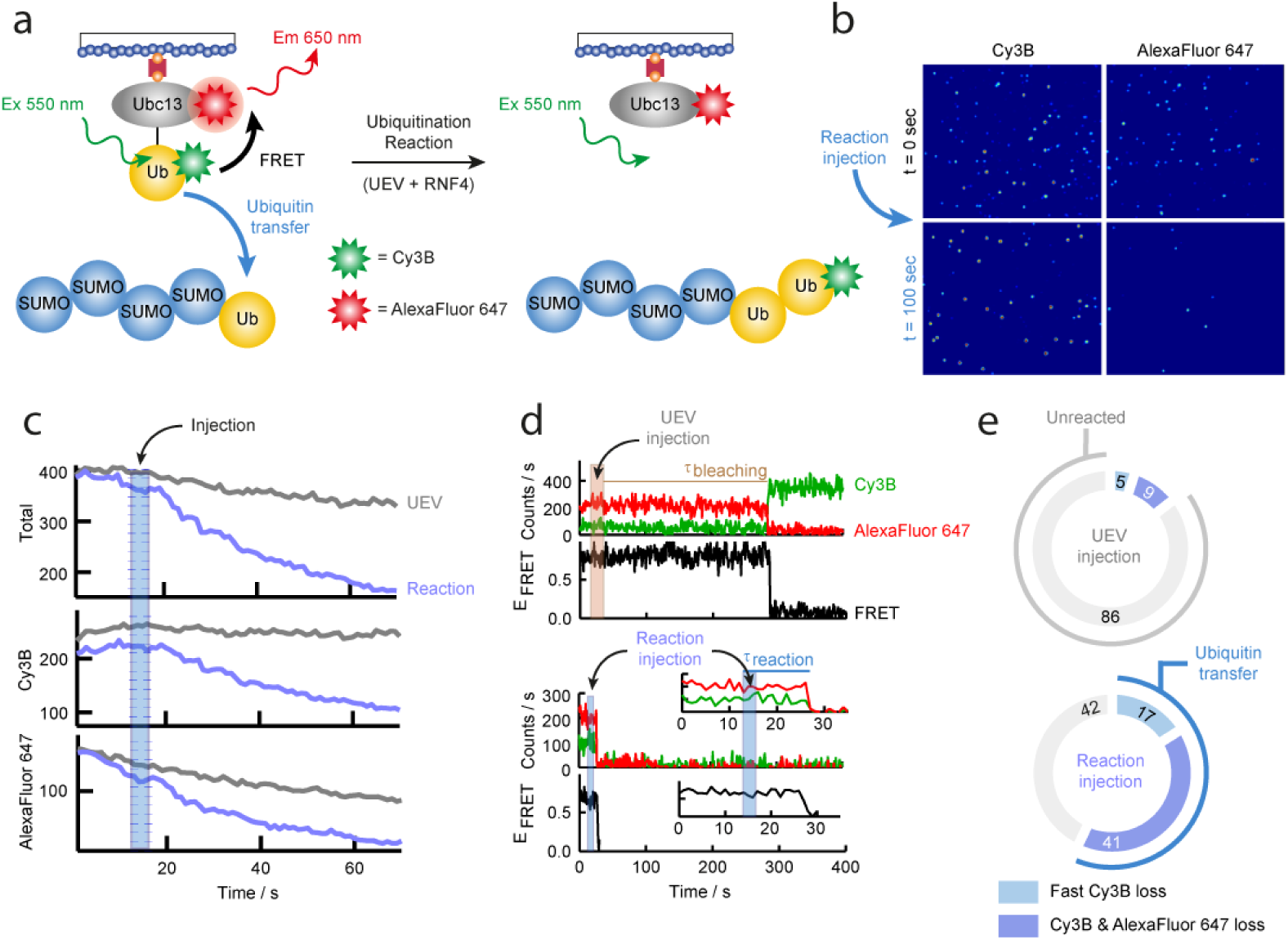
Ubiquitination reaction observed by real-time smFRET. **a**, Schematic diagram showing real-time ubiquitination reaction results in loss of FRET between the Ubc13~Ub conjugate and transfer of Cy3B labelled ubiquitin on to the substrate that diffuses into solution. **b**, Images of Cy3B and AlexaFluor 647 channels taken before (t = 0 seconds) and after (t = 100 seconds) the injection of reactants showing loss of both the Cy3B signal and the FRET signal from AlexaFluor 647 during the real-time smFRET experiment. **c**, Comparison of cumulative intensity variation from all fluorescent spots over time for UEV only injection (grey) versus reaction injection (blue). The total fluorescence intensity is shown along with separate Cy3B and AlexaFluor 647 intensities, with the injection interval highlighted in blue. **d**, Comparison of representative single molecule traces for the UEV injection (top) and reaction injection (bottom). Each molecule contains a Cy3B, AlexaFluor 647 and FRET intensity trace, with injection intervals highlighted in brown for UEV and blue for the reaction. The inset shows the first 35 seconds of the reaction injection, highlighting simultaneous loss and short lifetimes of the Cy3B, AlexaFluor 647 and FRET signals. **e**, Charts showing the percentage contribution of unreacted molecules and molecules undergoing ubiquitin transfer to the overall population, representing n = 263 and 270 individual molecules for the UEV and reaction injection respectively.

Contour-plots of FRET trajectories in a 90 second time-window following initiation of the ubiquitination reaction, showed rate of FRET loss due to ubiquitin transfer to substrate depends on RNF4 concentration and FRET loss was not observed in the absence of RNF4 or with an RNF4 mutant unable to bind the Ubc13~Ub conjugate (Fig. 4a). Single molecule analysis demonstrated that ubiquitin transfer rates can be distinguished from slow Cy3B and AlexaFluor 647 photobleaching events and quantified via dwell time of the FRET signal (Fig. 4b-c). The smFRET population histogram derived from only those molecules undergoing ubiquitin transfer, accurately shows the reaction proceeded from the high FRET state (E_FRET_~0.71) similar to that assigned to the closed conformation of the isopeptide linked E2~Ub conjugate (Fig. 4d, e). A low fraction of molecules reacting at a slower rate and from a low FRET state were also present in the UEV only injection, indicating that this was a background reaction that is not dependent on RNF4 (Fig. 4d-f).

**Figure 4.**
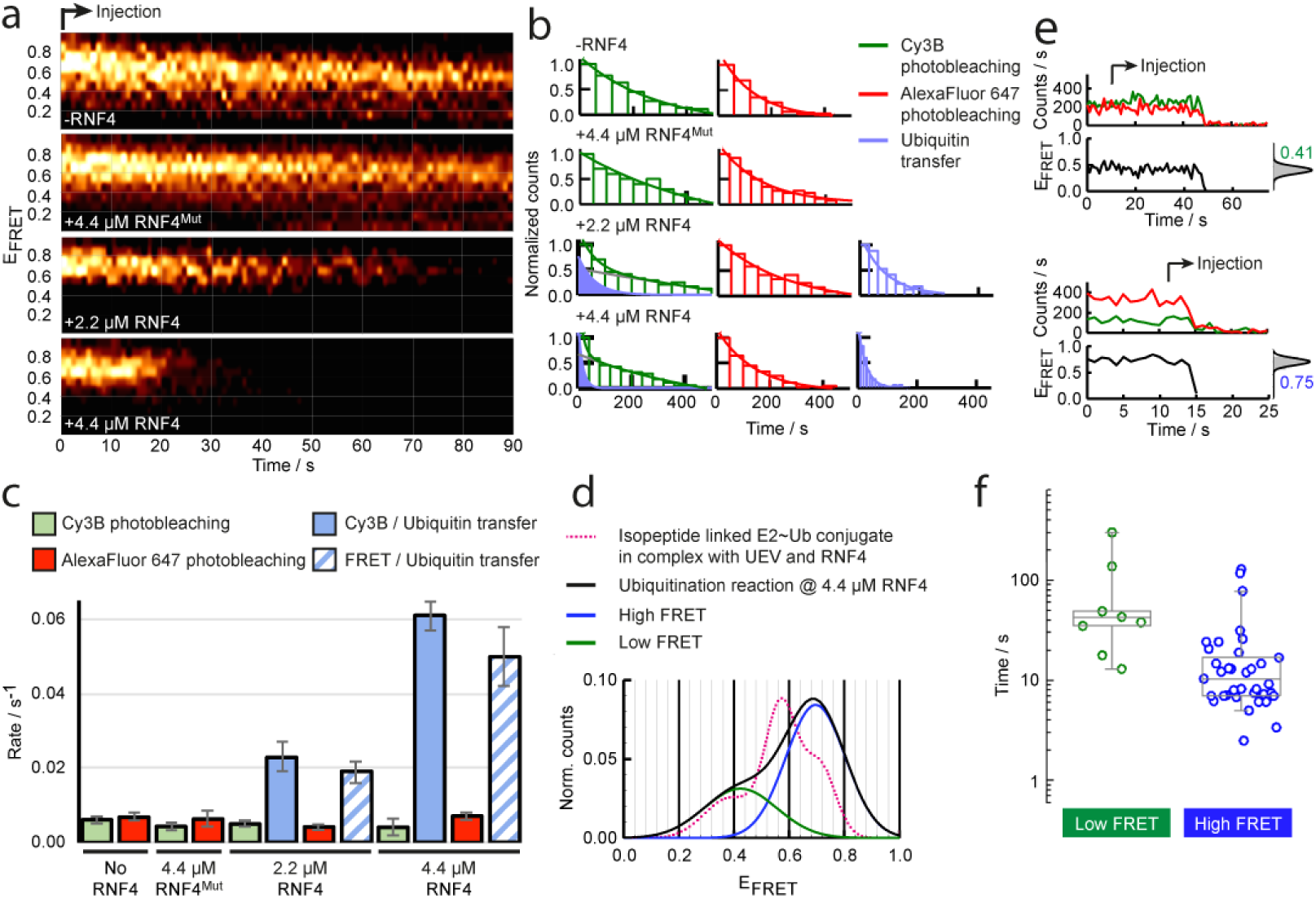
RNF4 catalyses the transfer of ubiquitin onto the substrate from a closed conformation of the Ubc13~Ub conjugate. **a**, Single molecule contour plots showing the progression of the FRET trajectory following the real-time injection of reactants at the indicated experimental conditions and concentrations. Mut denotes the RNF4 RING domain dimer containing E2~Ub binding site mutations (M140A and R181A) **b**, smFRET dwell-time histograms obtained for Cy3B photobleaching (green, left panel), fast Cy3B loss (blue, left panel), Alexa647 photobleaching (red, middle panel) and FRET loss events (blue, right panel). The solid lines represent the fit to single exponential decay functions. **c**, Comparison of the rates of Cy3B and AlexaFluor 647 photobleaching and the ubiquitination reaction, obtained by fitting the dwell-time histograms in **b** to mono- or biexponential decay functions. Error bars represent the standard error of the rate extracted from non-linear least squares fitting of the histograms in **b**. The ubiquitination reaction is split into the rate of loss of the FRET signal and the loss of Cy3B signal from molecules containing Cy3B only. **d**, Gaussian profile of relative FRET populations for the stable isopeptide linked conjugate and the real-time ubiquitination reaction. The histograms were omitted for clarity. **e**, Representative traces for reactions from low (top) and high (bottom) FRET states. Injection start time is indicated along with corresponding FRET histogram. **f**, Box-plot comparison of rate of reaction at 4.4 μM RNF4 from the low and high FRET states. The extremes, upper and lower quartiles of the distribution, and the median are represented by the whiskers, box and middle lines, respectively.

Previous crystallographic studies of RING E3 ligases bound to substrates and E2~Ub conjugates, show the conjugate restrained in the closed conformation with the substrate lysine aligned to attack the thioester bond^7,10,11^. Although yielding high-resolution structural information, these structures represent a snapshot of the arrangement of proteins prior to the catalytic step. NMR studies^5,15,17,18^ provide structural information on the conformational states of the E2~Ub conjugate, but such ensemble measurements require hours of averaging to generate a model. Stopped flow biochemical analysis^19^ generates rapid kinetics, but it lacks structural information. In this study, smFRET provides access to the conformation of individual E2~Ub conjugates in real-time undergoing rapid substrate ubiquitination from a closed conformation in a RING E3 ligase dependent reaction. Other E2~Ubls containing SUMO and Nedd8^10,11^ are similarly restrained in the closed conformation by their cognate RING E3 ligases, indicating that reaction from the closed conformation is likely to be a universal mechanism for RING E3 catalysis.

## Supporting information

All supplementary data

## Acknowledgements

We thank Anna Plechanovová and Gorjan Stojanovsky in the initial stage of the project. We thank Ulrich Zachariae of the University of Dundee for useful discussion. We thank the Division of Signal Transduction Therapy, University of Dundee for the gift of His-UBE1. This work was supported by grants from Cancer Research UK (C434/A21747), the Wellcome Trust (098391/Z/12/Z) to R.T.H.. J.C.P. thanks the University of St Andrews for financial support. R.T.H. is supported as a Senior Investigator of the Wellcome Trust.

## Author Contributions

E.B. cloned, expressed and purified proteins, conducted biochemical and smFRET experiments and interpreted data. J.C.P. conducted smFRET experiments and interpreted data. E.B. and J.C.P. contributed to data analysis. E.B., J.C.P. and R.T.H. wrote the paper. R.T.H. conceived the project and contributed to data analysis.

## Data Availability

All data are available from the authors upon request.

## Methods

### Cloning, expression and purification of recombinant proteins

Expression and purification of RNF4 and linear fusion RNF4 constructs was described previously^20^. Ub~4xSUMO-2 was also expressed and purified as described previously^21^. Human Ubc13 (also known as Ube2N) was previously subcloned into the pHISTEV30a vector using NcoI and HindIII restriction sites^7^. A sequence encoding an avitag (Gly-Leu-Asn-Asp-Ile-Phe-Glu-Ala-Gln-Lys-Ile-Glu-Trp-His-Glu) followed by a linker (Gly-Gly-Ala) was inserted between the TEV cleavage site and Ubc13 using the NcoI restriction site. A K24C mutation was introduced into Ubc13 using site-directed mutagenesis. Additional C87K and K92A mutations in Ubc13 were used as described previously^7^ to generate the stable isopeptide linked Ubc13~Ub conjugate. Ubc13 and Ube2V2 variants were expressed and purified as described previously^7^. As a result of cloning, Ubc13 has four extra residues (Gly-Ala-Met-Ser) before the N-terminal avitag after cleavage with TEV protease. His_6_ tagged Ubiquitin M-2C was expressed and purified as described previously^3,22^. An additional K63R mutation in ubiquitin was used to generate the unstable thioester linked Ubc13~Ub conjugate.

### Preparation of biotinylated and dye labelled protein

Avitagged Ubc13 was biotinylated using an enzyme called BirA as previously described^23^. The following reaction components were mixed together and incubated at 20 °C for 4 hours followed by 4 °C overnight: 10 mM Tris pH 7.5, 5 mM MgCl_2_, 200 mM KCl, 2.5 mM ATP, 0.5 mM d-biotin, 100 μM avitagged Ubc13 and 16 μM BirA. The reaction mixture was then purified by size exclusion chromatography with a HiLoad Superdex 75 16/600 column (GE Healthcare) and 50 mM Tris, 150 mM NaCl, and 0.5 mM TCEP, pH 7.5 as running buffer. Biotinylated Ubc13 K24C C87K K92A was buffer exchanged into 50 mM Tris, 150 mM NaCl, pH 7.0 using a Centri Pure Zetadex-25 gel filtration column (Generon) and labelled with Cy3B maleimide (GE Healthcare) at room temperature for 2 hours using a 5 times molar excess of Cy3B. Excess dye was removed using a Centri Pure Zetadex-25 gel filtration column and 50 mM Tris, 150 mM NaCl, and 0.5 mM TCEP, pH 7.5 as running buffer. His_6_ tagged Ubiquitin M-2C was also labelled using the same protocol but with Cy3B maleimide and AlexaFluor 647 C2 maleimide (Thermo Fisher Scientific) separately. All buffers were degassed and all dye labelling reactions and dye labelled proteins were protected from light.

### Preparation of isopeptide linked Ubc13~Ub conjugate

An isopeptide linkage was made between biotinylated and Cy3B labelled Ubc13 K24C C87K K92A and AlexaFluor 647 labelled his_6_ tagged Ubiquitin M-2C using a similar method as described previously^3,7^. The following mixture: 50 μM Ubc13, 60 μM Ubiquitin, 0.8 μM His_6_-Ube1, 3 mM ATP, 5 mM MgCl_2_, 50 mM Tris pH 10.0, 150 mM NaCl, 0.5 mM TCEP was incubated at 37 °C for about 21 hours. The reaction was then purified by Ni-NTA chromatography to isolate the isopeptide linked Ubc13~Ub conjugate and unreacted Ubiquitin, which are both his_6_ tagged. All buffers were degassed and all dye-labelled proteins were protected from light. Production and purification of the isopeptide linked Ubc13~Ub conjugate was analysed by SDS-PAGE and the FRET labelled protein was imaged using ChemiDoc MP (Bio-rad) with Cy3 and AlexaFluor 647 filter settings. Subsequent Coomassie blue staining allowed imaging of all proteins using the Coomassie filter setting. Ubiquitin I44A and Ubc13 L106A containing conjugates were prepared using the same protocol.

### Preparation of thioester linked Ubc13~Ub conjugate

A thioester linkage was made between biotinylated Ubc13 K24C and Cy3B labelled his_6_ tagged Ubiquitin M-2C K63R using the following mixture: 50 μM Ubc13, 50 μM Ubiquitin, 0.1 μM His_6_-Ube1, 3 mM ATP, 5 mM MgCl_2_, 50 mM Tris pH 7.5, 150 mM NaCl, 0.5 mM TCEP and 0.1 % NP-40 Alternative was incubated at 37 °C for 20 minutes. The reaction mixture containing the thioester product was buffer exchanged into 50 mM Tris, 150 mM NaCl, pH 7.0 using a Centri Pure Zetadex-25 gel filtration column. The thioester was labelled at biotinylated Ubc13 K24C with AlexaFluor 647 C2 maleimide at room temperature for 2 hours using a 5 times excess of AlexaFluor 647. Excess dye was removed using a Centri Pure Zetadex-25 gel filtration column and 50 mM Tris and 150 mM NaCl, pH 7.0 as running buffer. All buffers were degassed and all dye labelling reactions and dye labelled proteins were protected from light. Production and labelling of the thioester linked Ubc13~Ub conjugate was analysed by SDS-PAGE and the FRET labelled protein was imaged using ChemiDoc MP with Cy3 and AlexaFluor 647 filter settings. Subsequent Coomassie blue staining allowed imaging of all proteins in the samples using the Coomassie filter setting.

### Ubiquitination assays

The fluorescence polarization ubiquitination assay was performed and analysed as described previously using a fluorescein labelled version of Ubiquitin M-2C (Ub-5IAF)^22^. Briefly, 2.5 μM Ubc13, 2.5 μM UEV, 9.6 μM WT Ubiquitin, 0.4 μM Ub-5IAF, 0.1 μM His_6_-Ube1, 0.55 μM RNF4, 5.5 μM Ub~4xSUMO-2, 5 mM MgCl_2_, 50 mM Tris pH 7.5, 150 mM NaCl, 0.1% (v/v) NP-40 Alternative, 0.5 mM TCEP were mixed together and the reaction was incubated at 20 °C for 20 minutes taking samples at 0, 2, 5, 10, 15 and 20 minutes. 3 mM ATP was added after the 0 minute time point to start the reaction. The reaction was stopped at each time point by mixing it with stopping buffer (50 mM Tris pH 7.5, 150 mM NaCl, 0.5 mM TCEP, 0.1% (v/v) NP-40 Alternative, 150 mM EDTA pH 8.0) in a 2:1 ratio. Fluorescence polarization was measured using a PHERAstar FS microplate reader with 485 nm excitation and 520 nm emission wavelengths.

The substrate single turnover assay to validate labelled proteins was performed as previously described^20^. 15 μM FRET labelled thioester linked Ubc13~Ub conjugate was mixed in a 1:1 ratio with 15 μM UEV, 1.1 μM RNF4, 11 μM Ub~4xSUMO-2, 50 mM Tris pH 7.5, 150 mM NaCl, 0.1% (v/v) NP-40 Alternative, 0.5 mM TCEP and the reaction was incubated at 20 °C for 10 minutes and stopped with non-reducing SDS-PAGE loading buffer. Samples were taken at 0.5, 1, 2, 5 and 10 minutes. A WT Ubc13~Ub thioester linked conjugate, prepared using the same protocol as described above, was used as a control in this experiment. Ubiquitin in both Ubc13~Ub conjugates contained a K63R mutation such that only one ubiquitin was transferred onto the substrate. An RNF4 construct containing full length RNF4 linearly fused to a second RING domain was used in the ubiquitin reaction, similar to real-time smFRET experiments. Full length RNF4 includes four N-terminal SUMO Interaction Motifs, which engage the Ub~4xSUMO-2 substrate, and a C-terminal RING domain. One of the E2~Ub binding sites within the RING domain dimer was mutated (M140A and R181A) allowing only one E2~Ub conjugate to bind during the ubiquitination reaction. The assay was resolved by SDS-PAGE and the FRET labelled protein was imaged using ChemiDoc MP with Cy3 and AlexaFluor 647 filter settings. Subsequent Coomassie blue staining allowed imaging of all proteins in the assay using the Coomassie filter setting.

### Pull-down assay

Binding between MBP tagged RING domain dimer of RNF4 and the Ubc13~Ub conjugate was performed as described previously^20^. Assays were resolved by SDS-PAGE and results were analysed by western blotting with anti-ubiquitin primary antibody (Z0458, Dako) and anti-rabbit HRP secondary antibody for the unlabelled Ubc13~Ub conjugate. The FRET labelled Ubc13~Ub conjugate was analysed by SDS-PAGE and imaged using ChemiDoc MP with Cy3 and AlexaFluor 647 filter settings. Subsequent Coomassie blue staining allowed imaging of the proteins using the Coomassie filter setting.

### FRET dye modelling

Cy3 and AlexaFluor 647 were modelled on to the E2~Ub conjugate using the FRET-restrained positioning and screening (FPS) software^24^. As Cy3B maleimide is not available in the FPS software package, the closest alternative, Cy3 maleimide, was used for modelling purposes. Ubc13 (Chain B) and Ubiquitin (Chain C) in PDB 5AIT were used for modelling the accessible volume of FRET dyes as well as the distance between them for the closed conformation of the E2~Ub conjugate, while the NMR model (PDB accession number 2KJH) was used for the open conformation of the conjugate. As the cysteine in ubiquitin is contained within an N-terminal linker that is not present in these PDB files, the closest surface accessible amino acid, Gln2, was used for attachment and modelling the dye on ubiquitin. This is likely to result in a distance measurement that is shorter than expected; therefore, the resulting distance calculations were used as a guide.

### Slide passivation

Aminosilane treated slides were passivated with PEG-SVA and Biotin-PEG-SVA as described previously. Slides were assembled into four channels for stable isopeptide linked conjugate studies. Single channel slides with tubing and syringe attached for injection purposes were assembled for real-time reaction studies using the unstable thioester linked conjugate. Channels were coated with 0.2 mg ml^−1^ neutravidin for 10 minutes prior to addition of biotinylated FRET labelled proteins.

### Single molecule total internal reflection

smFRET experiments were performed as described previously^25^. All experiments were performed on a prism-type total-internal reflection microscope using an inverted microscope (Olympus IX71). A 532 nm laser (Crystalaser) was used for Cy3B excitation and images were collected on a back illuminated Ixon EMCCD camera (Andor, 512×512 pixels). Cy3B and AlexaFluor 647 fluorescence were split by dichroic mirrors (DCRLP645, Chroma Technology) into two channels allowing simultaneous imaging with Cy3B on the left and AlexaFluor 647 on the right of the EMCCD camera.

### Isopeptide linked conjugate smFRET experiments

50 pM of the stable isopeptide linked FRET labelled Ubc13~Ub conjugate was bound to a slide passivated with biotinylated PEG and neutravidin for 10 minutes. Excess free AlexaFluor 647 labelled ubiquitin was washed from the surface using 50 mM Tris, 150 mM NaCl, 0.5 mM TCEP, pH 7.5. Imaging buffer contained 50 mM Tris, 150 mM NaCl, 0.5 mM TCEP, pH 7.5, 1 mM Trolox, 6.25 mM 3,4-protocatechuic acid (PCA) and 250 nM protocatechuate dioxygenase (PCD). The Ubc13~Ub conjugate was imaged alone and in complex with UEV and the RING domain dimer of RNF4. One of the E2~Ub binding sites within the RING domain dimer was mutated (M140A and R181A) allowing only one E2~Ub conjugate to bind to the Ubc13~Ub conjugate. For complex imaging, imaging buffer was supplemented with 10 μM UEV and/or 40 μM RING domain dimer of RNF4. smFRET trajectories were acquired using 100 ms integration time.

### Thioester linked conjugate real-time smFRET experiments

A flow cell was generated as described previously^26^. Briefly, tubing was attached to the drilled holes at each end of the single channel slide. One piece of tubing was placed in an eppendorf containing the desired solution while a syringe was attached to the other piece of tubing through which the solution was drawn through the channel by suction. Using this suction method, 100 pM of the unstable thioester linked FRET labelled Ubc13~Ub conjugate was bound to a slide passivated with biotinylated PEG and neutravidin for 10 minutes. Excess free Cy3B labelled ubiquitin was washed from the surface using 50 mM Tris, 150 mM NaCl, pH 7.0. Free AlexaFluor 647 labelled Ubc13 is not observed due to direct excitation of Cy3B. Prior to the reaction the slide was washed with 50 mM Tris, 150 mM NaCl, 0.5 mM TCEP, pH 7.5. The UEV injection was performed with 2.5 μM UEV, 50 mM Tris, 150 mM NaCl, 0.5 mM TCEP, pH 7.5, 0.8% (w/v) d-glucose, 1 mM Trolox, 0.1 mg ml^−1^ glucose oxidase (Sigma) and 0.02 mg ml^−1^ glucose catalase (Sigma). The ubiquitination reaction injection was performed with 2.5 μM UEV, 2.2 - 4.4 μM RNF4 and 5.5 μM Ub~4xSUMO-2, 50 mM Tris, 150 mM NaCl, 0.5 mM TCEP, pH 7.5, 0.8% (w/v) d-glucose, 1 mM Trolox, 0.1 mg ml^−1^ glucose oxidase (Sigma) and 0.02 mg ml^−1^ glucose catalase (Sigma). An RNF4 construct containing full length RNF4 linearly fused to a second RING domain was used in the ubiquitin reaction, similar to the substrate single turnover ubiquitination assay used to validate the proteins. One of the E2~Ub binding sites within the RING domain dimer was mutated (M140A and R181A) allowing only one E2~Ub conjugate to bind during the ubiquitination reaction. Ubiquitin within the Ubc13~Ub conjugate contains a K63R mutation such that only a single ubiquitin is transferred onto a substrate molecule, rather than forming ubiquitin chains. smFRET trajectories were acquired with 1 s integration time.

### Data processing

IDL 6.0 was used to process an output containing intensity versus time for each single molecule trajectory detected by the EMCCD camera. Data were analysed in Matlab as described previously^27^. FRET efficiency (E_FRET_) was calculated from the raw Cy3B and AlexaFluor 647 intensity traces using *E*_*FRET*_ = *I*_*A*_/(*I*_*D*_ *+* α*I*_*A*_), where I_D_ = donor (Cy3B) intensity, I_A_ = acceptor (AlexaFluor 647) intensity and α = 0.88 accounts for 12% leakage into the AlexaFluor 647 detection channel. Histograms were prepared by averaging the first ten frames of each single molecule trace separately and these averages were combined to produce the overall FRET population. Histograms were normalized to compare percentage populations between samples.

