## Supplementary material for "Ubiquitin transfer by a RING E3 ligase occurs from a closed E2~Ub conformation": All supplementary data

**Extended Data Figure 1**

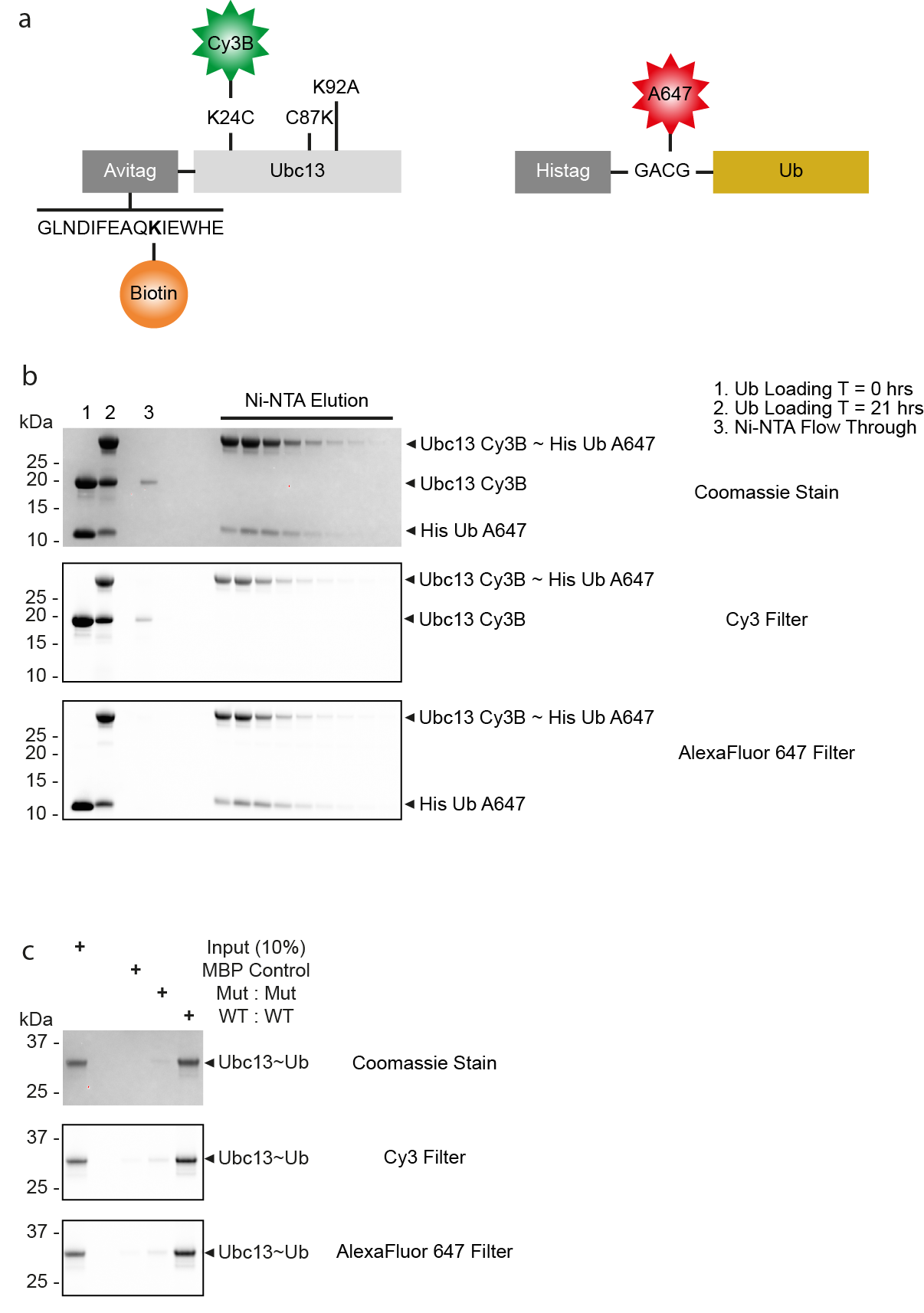

**Extended Data Figure 1 | Production, purification and validation of an isopeptide linked and FRET labelled Ubc13~Ub conjugate.**

**a**, Schematic diagrams of the Ubc13 and ubiquitin constructs used in FRET analysis of a stable isopeptide linked Ubc13~Ub conjugate. The Ubc13 construct contains a biotinylated N-terminal avitag followed by a short linker and Ubc13, which contains the mutations K24C, C87K and K92A. Ubc13 is labelled with Cy3B at position K24C. The ubiquitin construct contains an N-terminal histag and a short linker containing a cysteine for labelling with AlexaFluor 647, followed by ubiquitin. **b**, SDS-PAGE analysis showing generation of the stable isopeptide linked Ubc13~Ub conjugate and purification by Ni-NTA chromatography. **c**, RNF4 RING domain dimer pull-down experiment showing binding to the FRET labelled isopeptide linked Ubc13~Ub conjugate, analysed by SDS-PAGE. WT denotes a wild-type RING domain, while Mut denotes a RING domain containing E2~Ub binding site mutations (M140A and R181A).

**Extended Data Figure 2**

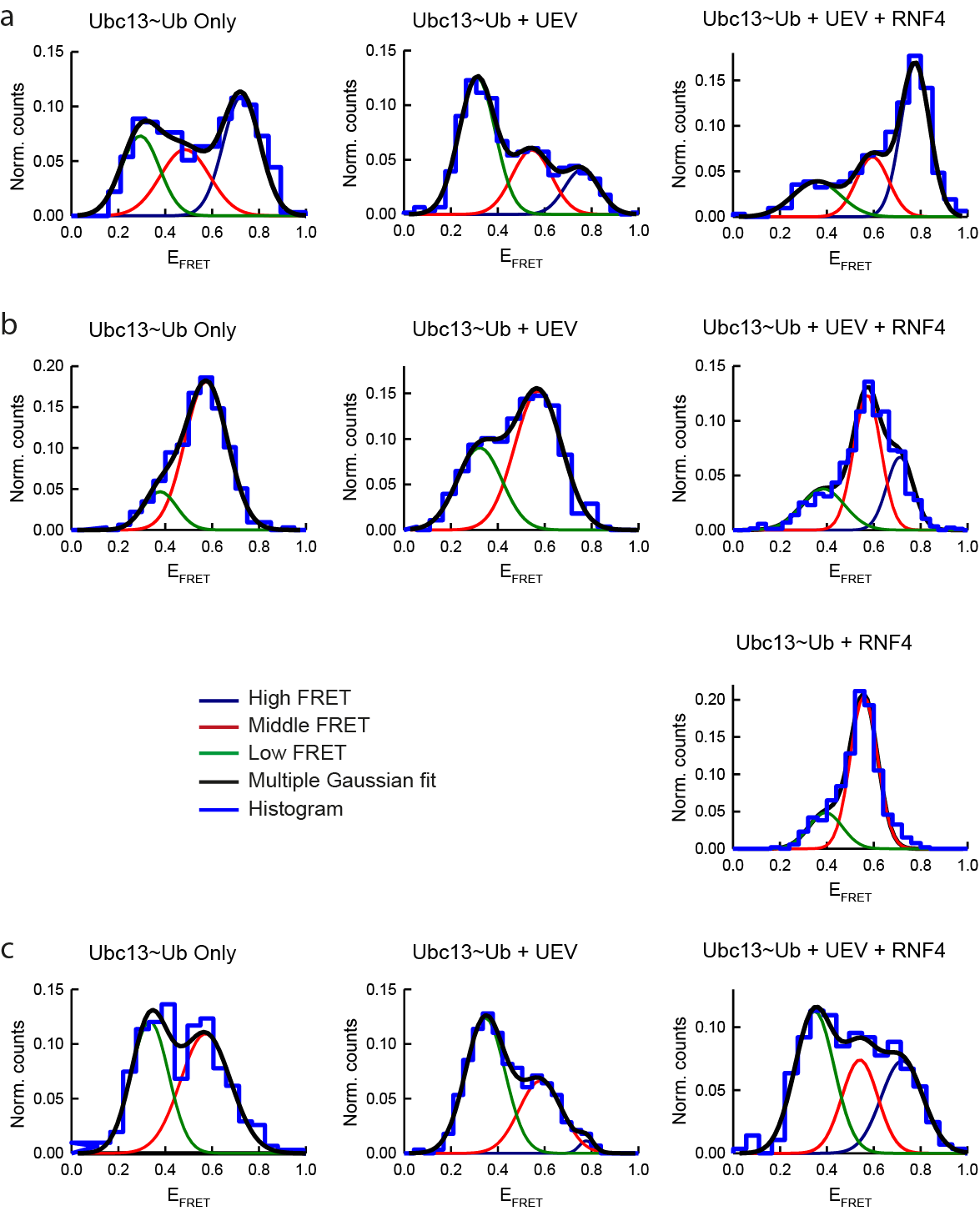

**Extended Data Figure 2 | smFRET histograms.**

**a**, Normalized smFRET histograms showing the E2~Ub conjugate conformation alone and in complex with UEV and the RNF4 RING domain dimer at 12 °C. **b**, Same as **a** but at 22 °C. The E2~Ub conjugate conformation in the presence of the RNF4 RING domain dimer only is also shown at 22 °C. **c**, Same as **a** but at 35 °C. For **a**, **b** and **c** the histogram is outlined in blue.

**Extended Data Figure 3**

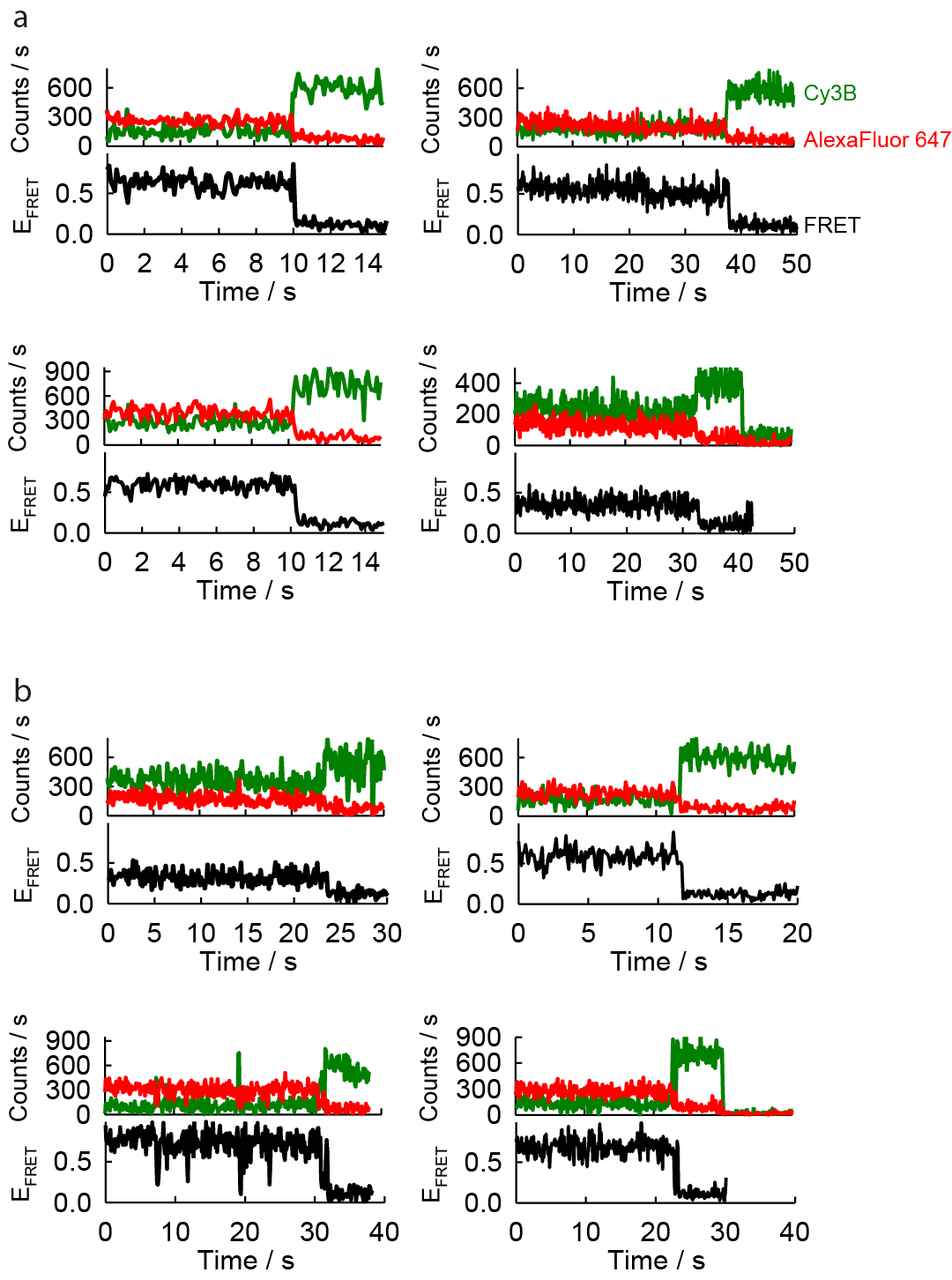

**Extended Data Figure 3 | Representative single molecule traces.**

**a**, Representative single molecule traces for the E2~Ub conjugate alone. Each molecule contains a Cy3B, AlexaFluor 647 and FRET intensity trace. **b**, Same as in **a** but for the E2~Ub conjugate in complex with UEV and RNF4 RING domain dimer.

**Extended Data Figure 4**

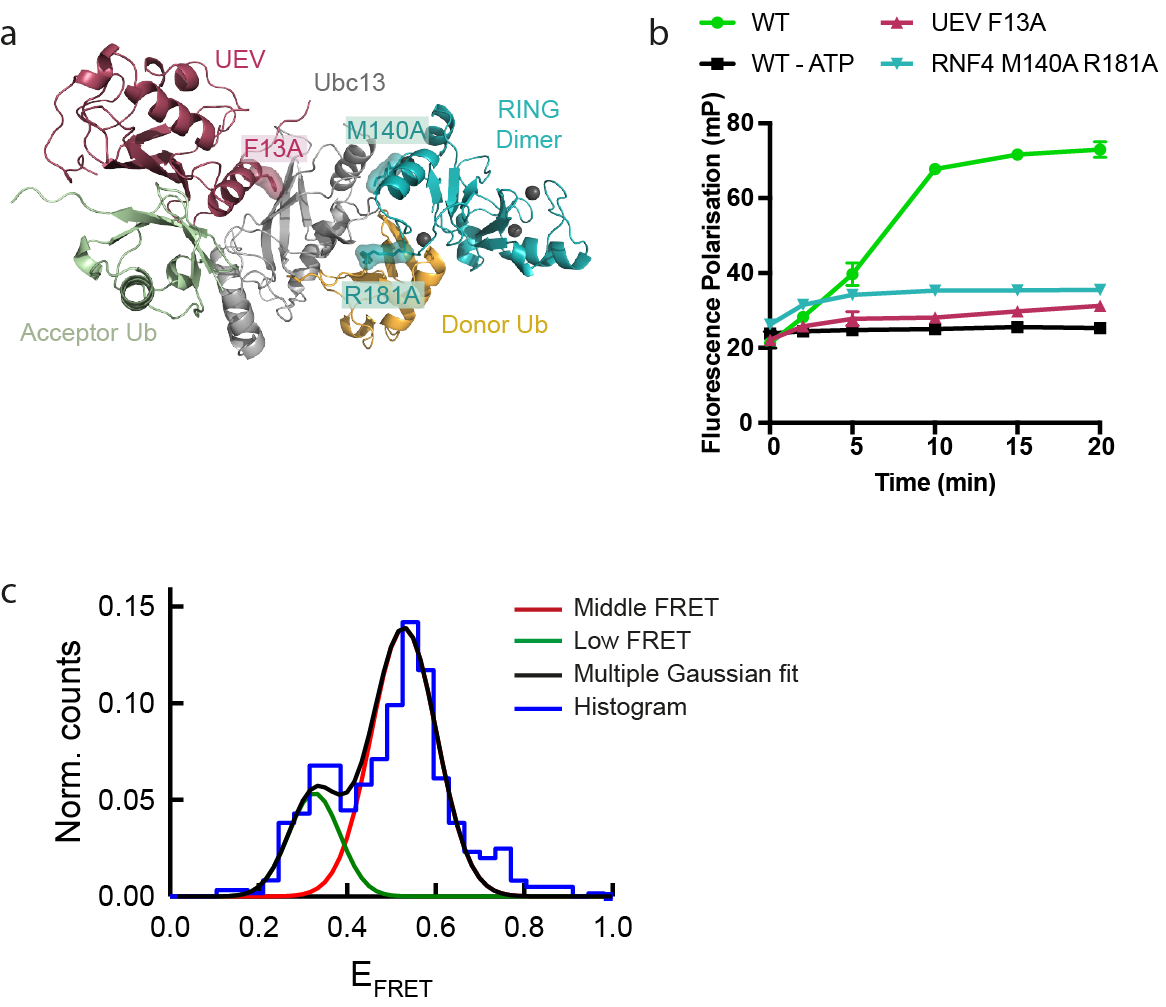

**Extended Data Figure 4 | Effect of negative control complex mutations on the conformational state of the E2~Ub conjugate.**

**a**, Locations of mutations made within the complex (PDB accession number 5AIT). **b**, Fluorescence polarization ubiquitination assay showing the effect of mutations on K63-linked polyubiquitin chain formation, represented as mean ± s.d. of triplicate measurements. Error bars are omitted when the error is smaller than the data point. **c**, Normalized smFRET histogram showing the E2~Ub conjugate conformation in the presence of UEV F13A and the RNF4 RING domain dimer containing M140A and R181A mutations. The histogram is outlined in blue.

**Extended Data Figure 5**

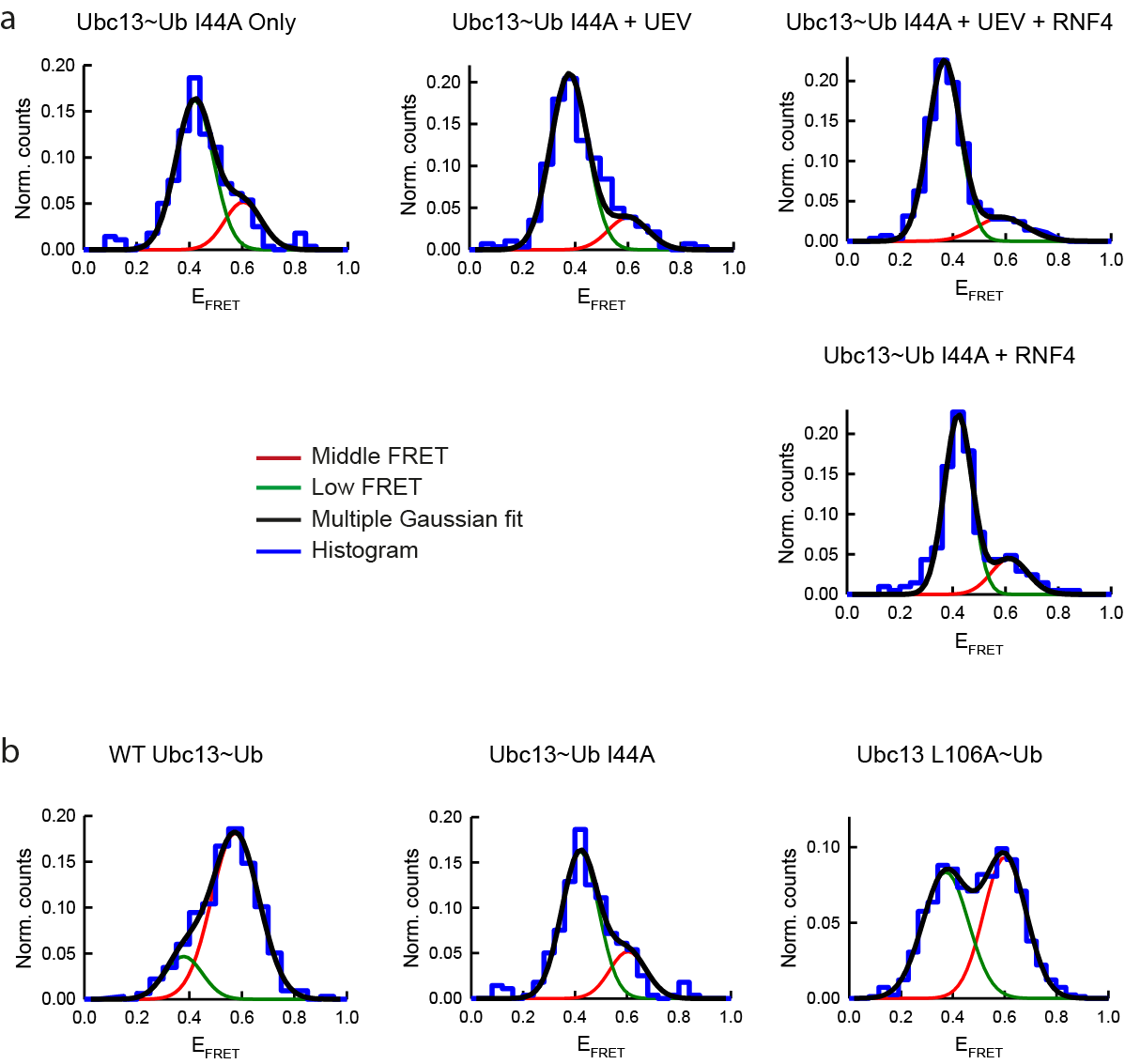

**Extended Data Figure 5 | smFRET histograms for Ub I44A and Ubc13 L106A containing Ubc13~Ub conjugates.**

**a**, Normalized smFRET histograms showing the E2~Ub conjugate conformation containing Ub I44A alone and in complex with UEV and the RNF4 RING domain dimer and in the presence of the RNF4 RING domain dimer only. **b**, Normalized smFRET histograms showing the conformation of the WT E2~Ub conjugate and the E2~Ub conjugate containing either Ub I44A or Ubc13 L106A. For **a** and **b**, the histogram is outlined in blue.

**Extended Data Figure 6**

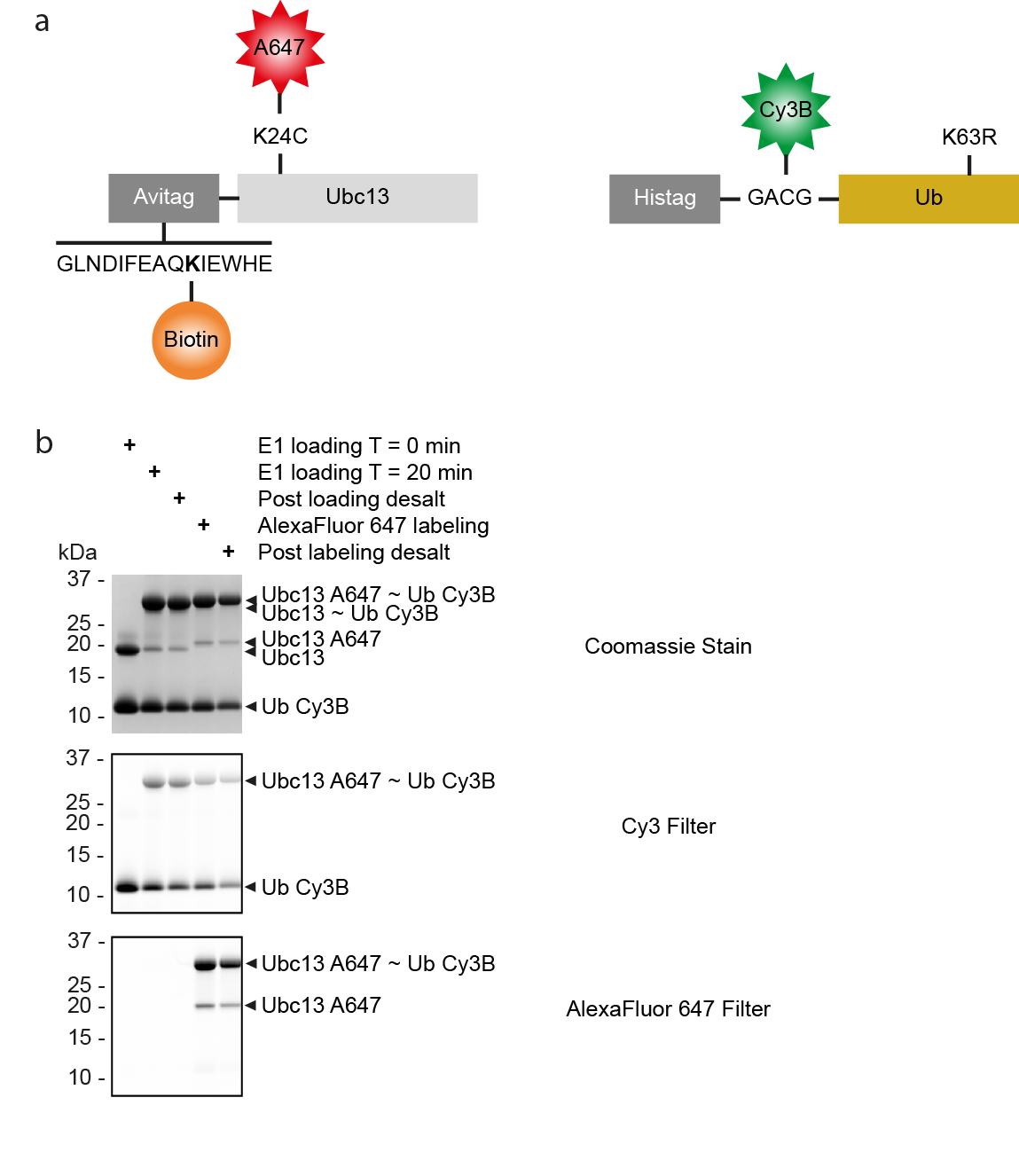

**Extended Data Figure 6 | Production and labelling of a thioester linked and FRET labelled Ubc13~Ub conjugate.**

**a**, Schematic diagrams of the Ubc13 and ubiquitin constructs used in FRET analysis of an unstable thioester linked Ubc13~Ub conjugate. The Ubc13 construct contains a biotinylated N-terminal avitag followed by a short linker and Ubc13. Ubc13 contains the mutation K24C that is used for labelling with AlexaFluor 647. The ubiquitin construct contains an N-terminal histag and a short linker containing a cysteine for labelling with Cy3B. This is followed by ubiquitin containing a K63R mutation. **b**, SDS-PAGE analysis showing generation of the unstable thioester linked Ubc13~Ub conjugate and labelling with AlexaFluor 647.

**Extended Data Figure 7**

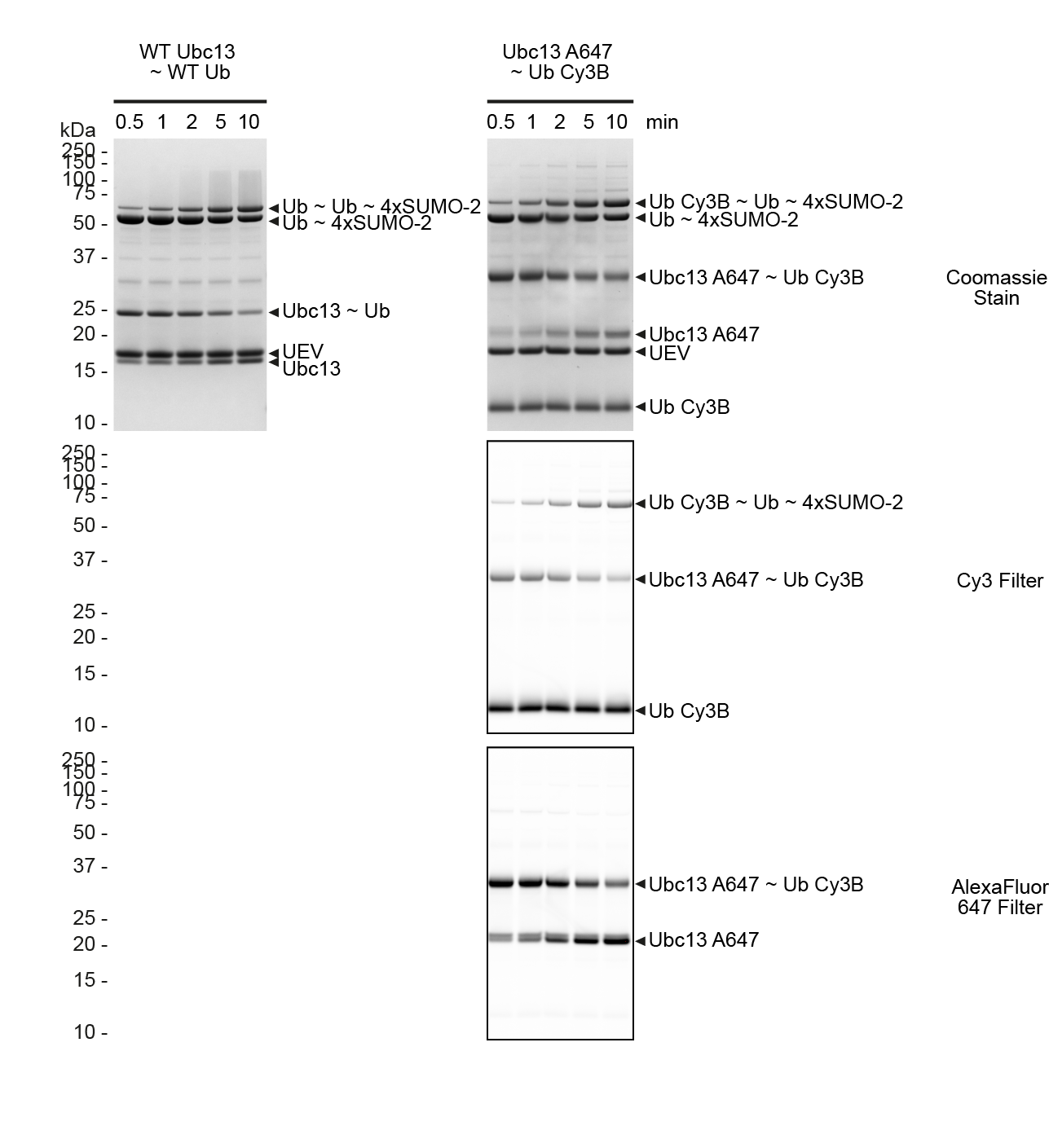

**Extended Data Figure 7 | Validation of a thioester linked and FRET labelled Ubc13~Ub conjugate.**

SDS-PAGE analysis of a substrate single turnover ubiquitination assay showing the ability of a WT Ubc13~Ub compared to the tagged and FRET labelled Ubc13~Ub conjugate to monoubiquitinate the Ub~4xSUMO-2 substrate.
